# Specialization and selective social attention establishes the balance between individual and social learning

**DOI:** 10.1101/2021.02.03.429553

**Authors:** Charley M. Wu, Mark K. Ho, Benjamin Kahl, Christina Leuker, Björn Meder, Ralf H.J.M. Kurvers

## Abstract

A key question individuals face in any social learning environment is when to innovate alone and when to imitate others. Previous simulation results have found that the best performing groups exhibit an intermediate balance, yet it is still largely unknown how individuals collectively negotiate this balance. We use an immersive collective foraging experiment, implemented in the Minecraft game engine, facilitating unprecedented access to spatial trajectories and visual field data. The virtual environment imposes a limited field of view, creating a natural trade-off between allocating visual attention towards individual search or to look towards peers for social imitation. By analyzing foraging patterns, social interactions (visual and spatial), and social influence, we shine new light on how groups collectively adapt to the fluctuating demands of the environment through specialization and selective imitation, rather than homogeneity and indiscriminate copying of others.

## Introduction

Most of what we learn, we learn from other people. Social learning often provides a cheap alternative to individual trial and error learning (Kendal et al., 2018), whereby observing and imitating the successful actions of others provides an expedient pathway to rewards. However, social learning can easily go awry: too many social learners in a population can create maladaptive information cascades (Toyokawa, Whalen, & Laland, 2019; Tump, Pleskac, & Kurvers, 2020) of imitators imitating other imitators, with poor outcomes at both the individual and collective level (Rogers, 1988).

Deciding when to learn socially or individually is a strategy selection problem (Payne, Bettman, & Johnson, 1988). Yet in contrast to previous work in purely individual domains (Marewski & Schooler, 2011; Rieskamp & Otto, 2006), arbitrating between social and individual learning is complicated by *frequency-dependent fitness*: one’s performance depends on the frequency of strategies used by other individuals in the population (Laland, 2004; Rogers, 1988). Thus the dominant strategy depends on what strategy other people choose: social learning is profitable when rare, but fails amidst a glut of imitators.

The trade-off between individual and social learning has been well-studied in socially foraging animals (but less so in humans) using the *producer-scrounger* game (Barnard & Sibly, 1981; Kurvers et al., 2009). A group of animals are typically placed in an environment with spatially distributed reward patches, presenting a type of game theory dilemma: playing the “producer” strategy is to use individual learning to discover new reward patches, while playing the “scrounger” strategy is to imitate others by foraging from patches they have discovered. Because of the frequency-dependent nature of scrounging, it is expected that over repeated interactions, individuals in a group will theoretically gravitate towards a mixed equilibrium of individuals who engage predominately in producing and others who focus on scrounging, yielding an intermediate balance of individual and social learning in the population (Henrich & Boyd, 1998; Ehn & Laland, 2012; Rogers, 1988). In line with these predictions, human participants have been shown to assort themselves into balanced mixtures of individual and social learners (Kameda & Nakanishi, 2002).

While people are certainly capable of flexible arbitration rather than itinerant deployment of a fixed strategy (Miu, Gulley, Laland, & Rendell, 2020; Kendal et al., 2018), we lack a clear descriptive understanding of the mechanisms that allow individuals to adapt to changes in both the asocial and social environment. The predictability of rewards in the asocial environment impacts the relative value of social learning over individual learning, which can change the optimum ratio of strategies in the population. At the same time, the frequency of social learning in a population presents a dynamically changing social environment, altering the optimal individual strategy at a given point in time.

Previous work studying arbitration between individual and social learning has typically used problems where the asocial and social environments are static (e.g., bandit or lottery tasks with advice from a fixed agent; Zhang & Gläscher, 2020; Diaconescu et al., 2020). However, in naturalistic human interactions, the asocial reward environment is dynamic, as individuals compete over finite resources and resource deplete over time. The social environment is also dynamic as peers adapt their learning strategies, and individuals create their own social interaction networks. Here we investigate the flexible and dynamic arbitration between individual and social learning in conditions where both the asocial and social environments dynamically and naturally fluctuate over time.

### Goals and scope

Our focus is two-fold. First, we want to understand meta-cognitive control between social and individual learning strategies at the individual level, and test whether people can flexibly adapt their strategy use to different reward environments that alter the relative value of social learning. For this, we rely on visual field data, which provides a marker of social attention directed towards peers. Second, we want to understand the mechanisms that allow individuals in groups to dynamically negotiate a balance between social and individual learning at the collective level, where we deploy network analyses and measure social influence through “pull events” in foraging trajectories.

We address these questions using a collective search task implemented in an immersive virtual environment through the Minecraft game engine (Fig 1a). The task is modeled on the producer-scrounger game, where a limited field of view creates an attention allocation problem: visual attention can either be allocated to focus on individual exploration or to look towards peers for social imitation. Employing high-resolution spatial tracking and automated transcription of visual field data, we provide an unprecedented level of detail into the interaction dynamics that negotiate the balance between individual and social learning.

**Figure 1:**
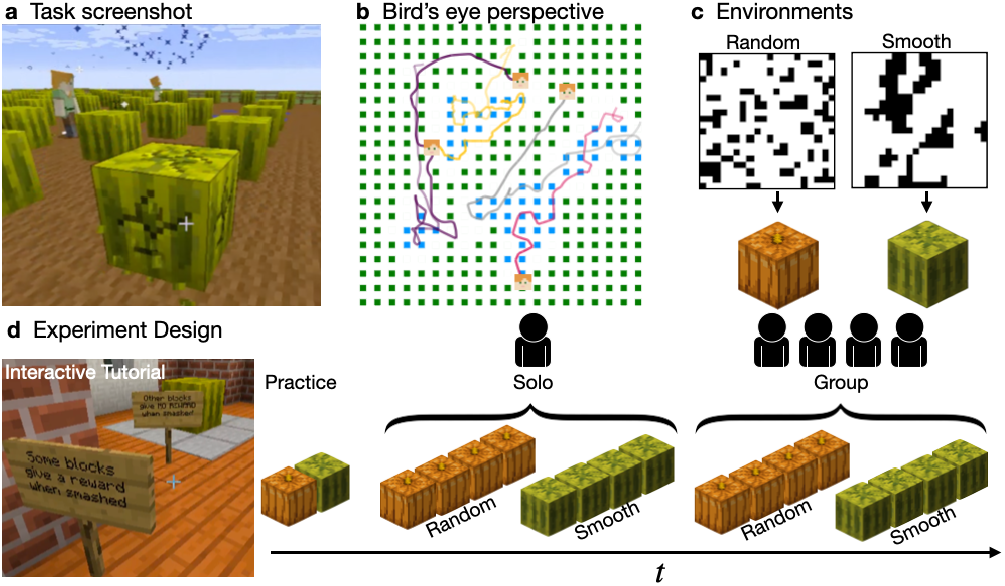
Experiment. **a**) Screenshot (cropped) from a social round. A reward discovered by another player is visible as a blue splash in the air. **b**) A bird’s eye depiction of the task, showing the full 60×60 field and participant trajectories over a two minute round. Blue blocks correspond to discovered rewards, while gaps in the regular grid of green blocks indicates no reward. **c**) Examples of smooth and random environments, which were mapped to either melons or pumpkins (counterbalanced across sessions). Each pixel indicates either reward (black) or no reward (white). Gaps between blocks are omitted. **d**) Experiment design. Participants first completed an interactive tutorial (https://youtu.be/bj55n8CI_Nk), followed by a training round of each environment. The main task was a 2×2 within-participant design, manipulating reward environment (smooth vs. random) and search condition (solo vs. group). Each of the four configurations was completed in four consecutive rounds of the same type (16 in total). Round order was pseudo-randomized and counterbalanced between sessions.

Within subjects, we manipulate i) whether search is performed alone or in groups of four, and ii) the predictability of rewards in the environment, altering the relative effectiveness of social and individual learning (Barkoczi, Analytis, & Wu, 2016; Rendell et al., 2010). Between environments, we find that participants flexibly adapt their level of social learning, relying more on social information when profitable. In environments where predictable rewards favor social learning, participants use a combination of specialization and selective attention. Rather than all individuals homogeneously converging on the same intermediate strategy, participants become specialized as either the target or the source of social attention. This produces an asymmetric social network structure where those who specialize in individual learning are the most observed. Thus, social learning is selective towards strongly individual learners, rather than copying indiscriminately.

### Methods

#### Participants and materials

Participants (*N* = 44) were recruited^1^ from the recruitment pool of the Max Planck Institute for Human Development in Berlin (MPIB) and selected to be between the ages of 18 and 30 to minimize generational differences in exposure to first person computer games (mean age: 26.5 ± 4.6 SD; 25 Females, 1 non-binary). The study was approved by the MPIB Institutional Review Board (A 2019-05) and participants signed an informed consent form prior to participation. Participants earned a base payment of €12 plus a bonus of €0.03 per reward, earning on average €17.21 ± 0.88.

The experiment was conducted in a computer lab using a modified Minecraft server, where participants controlled an avatar from a first person perspective (Fig. 1a). Each round of the experiment took place on a 60×60 field (bounded by a fence), containing 400 resource blocks, laid out in a 20 × 20 grid with a two block gap between each block (Fig. 1b). Each resource block (either watermelon or pumpkin, depending on environment) could be destroyed by continually hitting it (holding down left mouse button) for 2.25 seconds, yielding a binary outcome of either reward or no reward.

Rewards were indicated by a blue splash effect (Fig. 1a), visible from any position on the map. Only resource blocks (watermelon or pumpkin) were capable of being destroyed in the experiment and were not renewed within the round. Blocks did not possess any *a priori* visual features indicating whether or not they contained a reward. However, rewards in smooth environments were partly predictable, since observing a reward predicts other rewards nearby (Fig. 1c). In contrast, rewards in random environments were drawn from a uniform distribution (without replacement) and thus not predictable. Participants were individually incentivized to collect as many rewards as possible, which were translated into a bonus payment at the end of the experiment.

#### Design and procedure

Participants completed the task in randomly assigned groups of four, with group members completing the task in the same room and made aware they were interacting with each other. After an in-game tutorial and two practice rounds (see below), participants completed 16 rounds of the task, each lasting two minutes. We used a 2×2 within-subject design, manipulating the reward environment (random vs. smooth) and search condition (solo vs. group), with each combination completed in four sequential rounds (Fig. 1d).

The reward environment of a given round (random vs. smooth) was made salient by the use of either pumpkin or watermelon blocks (counter-balanced across groups; Fig. 1c). Both environments had the same number of rewards (25% of blocks), but with rewards either randomly or smoothly distributed. Random environments were generated by uniformly sampling 25% of blocks (without replacement). Smooth environments were designed to contain clustered reward distributions, which varied smoothly over space. We first sampled a bivariate reward function from a Gaussian Process prior, where we used a radial basis function kernel with the lengthscale parameter set to 4 (similar to Wu, Schulz, Garvert, Meder, & Schuck, 2020). Sampled reward functions were then binarized, such that the top quartile (25%) of block locations were set to contain rewards. In the tutorial, participants were given verbal descriptions of each reward condition, saw two fully-revealed illustrations of each environment class from a bird’s-eye perspective (Fig. 1c), and interactively destroyed a 3 × 3 patch of both environments.

The search condition of a given round was made salient by having participants either stand on an isolated teleportation platform (solo) or on a common teleportation platform with the other participants (group) in order to start the round. In the solo condition, participants searched on identical replications of the same environments but without interactions with one another. In the group condition, participants searched on the same environment and could compete with and imitate one another.

After receiving verbal instructions, participants completed a tutorial to familiarize themselves with the controls, how to destroy blocks, the difference between smooth and random reward distributions, and the overall task structure (Fig. 1d). They then completed two practice rounds in both smooth and random environments, which were identical to the solo condition of the main task (but without contributing to bonus payments). Each round lasted two minutes, with the end of the round corresponding to the sun setting below the horizon. This served as an approximate in-game timer for each round, and was communicated to participants in the tutorial. A three second countdown timer was also shown on the screen. At the end of the round, participants were given an on-screen announcement indicating the number of rewards they earned and notifying them of the reward environment and search condition for the next round. Participants were then teleported into a lobby (either separate lobbies for solo or a communal lobby for group rounds), and were required to all stand on a “teleportation” block to indicate readiness for the next round. Then players were teleported into a random position in the next environment, facing a random direction. For the entire experiment the sound was turned off, participants could not see each other’s screens, and task-irrelevant controls (e.g., crafting menu) were disabled.

#### Data collection

Experimental data was collected using a custom data logging module programmed in Java, separated into player logs and map logs. Player logs contained each player’s [*x, y*] spatial positions together with the pitch and yaw components of their visual orientation sampled at 20hz (every 0.05s). Map logs contained information about each player’s interactions with resource blocks, also sampled at 20hz. Together, the player and map logs allowed us to completely reconstruct the visual fields from all participants using a simulation programmed in the Unity game engine. This allowed us to automatically transcribe visibility information. We simulated each player’s field of view with all entities and other players rendered using a unique RGB color mask. From each frame of the simulations, we analyzed the pixels of the resulting image to determine which players and entities were visible at a given time.

### Results

We first focus on task performance before analyzing social interactions using visual and spatial data. Lastly, we model social influence by detecting pull events in foraging patterns.

#### Task performance

Using a two-way within subjects ANOVA, we found that participants acquired more rewards in smooth environments (*F*(1, 43) = 154.0, η^2^ = .47, *p* < .001) and in the solo condition (*F*(1, 43) = 8.54, η^2^ = .02, *p* = .006), with an additional interaction of smooth:solo (*F*(1, 43) = 6.42, η^2^ = .01, *p* = .015; Fig. 2a). This improved performance in the smooth:solo condition was mediated by learning over rounds (Fig. 2b). We fit a Bayesian mixed effects regression using environment, search condition, and round number to predict reward, while treating participants as random effects. The only reliable interaction with round was found in the smooth:solo condition (*b* = 0.07, 95% Highest Posterior Density (HPD): [0.01,0.12], *p*(*b* > 0) = .99). Thus, participants improved their ability to detect rewards in smooth environments over successive rounds when searching alone, but not in groups.

**Figure 2:**
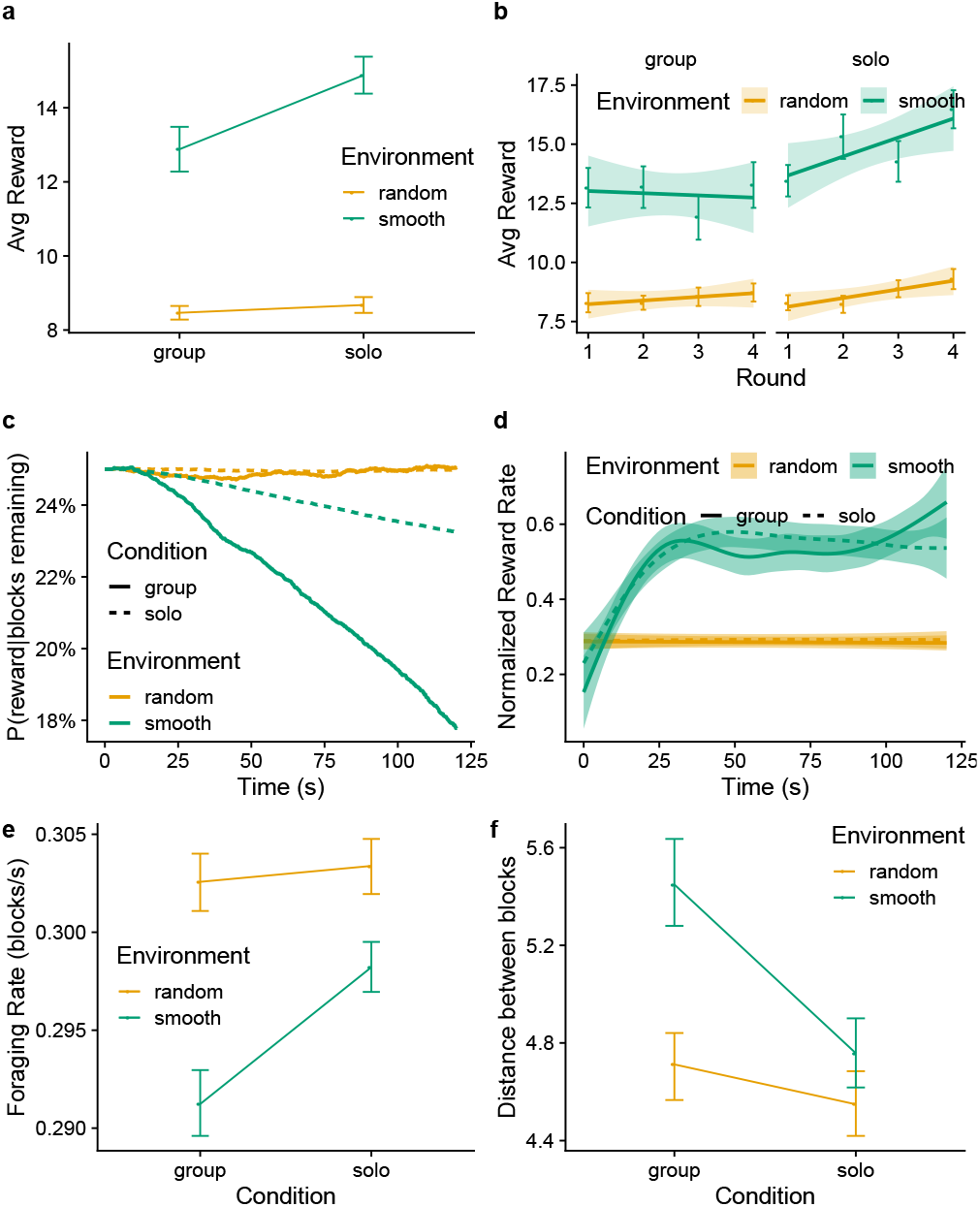
Behavioral results. **a**) Average reward across conditions. Dots indicate group means with error bars showing the standard error. **b**) Average over rounds, where each dot and error bar show the mean and standard error, and the lines and ribbons show the mean and 95% CI of a linear regression. **c**) The probability that a randomly sampled block contains a reward at any given time. This illustrates the higher depletion of rewards in more predictable environments (i.e., smooth) and when more participants are competing for the same finite number of resources (i.e., group). **d**) Normalized reward rate shows the average rate of rewards, normalized by the expectation of rewards (panel c). Lines and ribbons show the mean and 95% CI of a generalized additive model. **e**) Foraging rate defined in terms of blocks destroyed per second. **f**) The average distance between destroyed blocks.

However, these reward differences were substantially influenced by the dynamics of reward depletion (Fig. 2c), where more predictable rewards (i.e., smooth) and more participants searching for the same finite number of rewards (i.e., groups) both contributed to a faster decay in the baseline probability that one of the remaining blocks contained a reward. Thus, we computed the normalized reward rate (Fig. 2d), by normalizing the instantaneous reward rate by the current expected reward rate *P*(*reward|blocksRemaining*). This measure shows how well participants perform relative to a dynamically changing random baseline. We found higher performance in smooth environments (*F*(1, 43) = 182.5, η^2^ = .53, *p* < .001), but no difference between group or solo conditions (*F*(1, 43) = 0.33, η^2^ = .001, *p* = .57). We also no longer found any significant interaction in the group:smooth condition (*F*(1, 43) = 0.03, η^2^ < .001, *p* = .863). Thus, when accounting for differences in expected rewards due to depletion, participants performed equivalently in the solo and group conditions.

Participants’ foraging patterns were also influenced by the environment and social search conditions. Figure 2e shows the average foraging rate, which is defined as the number of blocks destroyed per second. Participants were more selective and foraged slower in smooth environments (*F*(1, 43) = 5.15, η^2^ = .02, *p* = .028) and in the group condition (*F*(1, 43) = 7.39, η^2^ = .01, *p* = .009). There was also an interaction between search condition and environment (*F*(1, 43) = 8.20, η^2^ = .003, *p* = .006), where participants were especially slower when combining the predictably smooth rewards with group dynamics.

These adaptive patterns of foraging selectivity are also present when analyzing the distance between destroyed blocks (Fig. 2f). Participants travelled further between foraged blocks in smooth environments (*F*(1, 43) = 14.3, η^2^ = .07, *p* < .001) and in the group condition (*F*(1, 43) = 34.6, *p* < .001, η^2^ = .07). We again see the same interaction between search condition and environment (*F*(1, 43) = 15.1, *p* < .001, η^2^ = .03), where participants in the group:smooth condition especially foraged over the longest distances.

#### Social Interactions

Next, we focus on the social interactions between participants in the group rounds. By recreating all experimental data in the Unity game engine (see methods), we were able to programmatically annotate all field of view (FOV) data. This allowed us to determine when any given participant was visible to any other participant at all points in time. We then computed the average number of visible peers as a proxy for *social attention* (Fig. 3a), which is higher when more peers were visible and for longer durations. Consistent with the fact that social information had no value in random environments (due to unpredictable rewards), we found higher social attention in smooth environments (paired *t*-test: *t*(43) = 2.7, *d* = 0.5, *p* = .011). There was also a marginal interaction effect of round, where social attention tended to decrease over rounds in the random environment (Bayesian mixed effects regression: *b* = −0.01, 95% CI: [−0.02, 0.01], *p*(*b* < 0) = .86; Fig. 3b). Thus, participants observed other participants less in random environments (where social information had no value), with the magnitude of this difference increasing over successive rounds.

**Figure 3:**
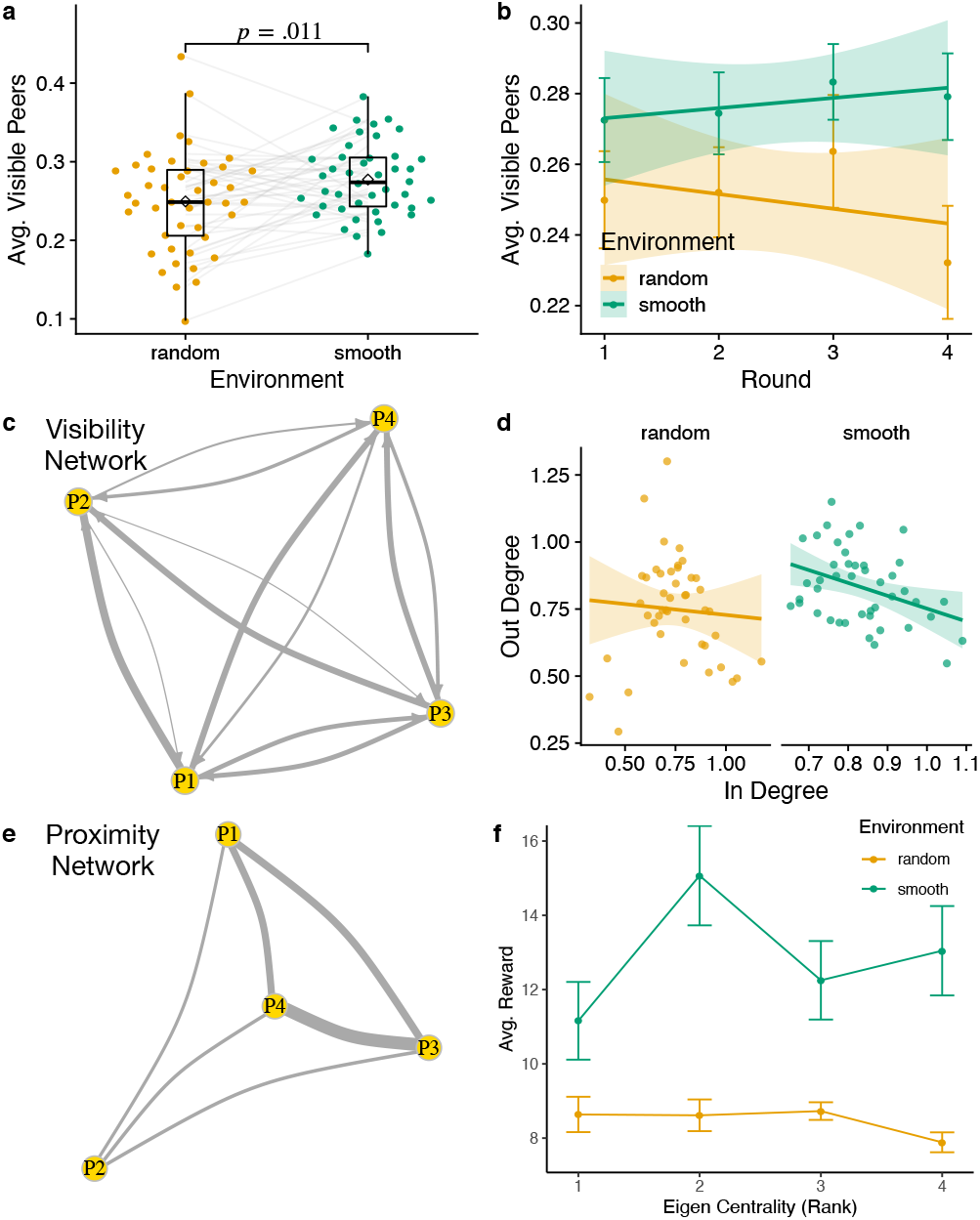
Social interaction results (group rounds only). **a**) The average number of visible peers (at any point in time). Each connected dot is a participant, with an overlaid Tukey boxplot providing group-level statistics. The diamond show the group means. The *p*-value is based on a paired *t*-test. **b**) Changes in social visibility over consecutive rounds. Dots and error bars indicate the aggregate mean and standard error, while the line and ribbon show the mean and 95% CI of a linear regression. **c**) An example of a visibility network, where each node is a participant and the directed edges are weighted by the proportion of time (within a round) that the target participant is visible to the observer. **d**) A comparison of the in- and out-degree of each participant (dot) with the lines indicating a linear regression. **e**) An example of a spatial network, where the undirected edges are weighted by the average spatial proximity between participants. **f**) Correspondence between the eigenvector centrality rank (how connected one is to other connected nodes) of each participant with their average reward. Dots and error bars show the aggregate mean and standard error.

In order to better understand the pairwise interactions between participants’ visual attention, we constructed *visibility networks* for each round (Fig. 3c), where the directed edges are weighted based on the amount of time one participant observed another. Based on these graphs, we computed the weighted in- and out-degree for each participant (Fig. 3d). Higher in-degree corresponds to being observed more frequently (i.e., “celebrity” factor), while higher out-degree corresponds to observing others more frequently (i.e., “paparazzi” factor). In smooth environments, we find a negative correlation between in- and out-degree (*r* = −.37, *p* = .014), but no correlation in random environments (*r* = −.07, *p* = .652). We obtain similar results using rank correlation (smooth: *r_τ_* = −.21, *p* = .044; random: *r_τ_* = −.11, *p* = .275). This suggests that participants were more specialized in smooth environments, with celebrities who were frequently observed but rarely observed others, and paparazzi who observed others intensely but were seldom observed themselves.

We also built *proximity networks* where the undirected edges were weighted by the spatial proximity between participants (stronger edge weights for closer average distance; Fig. 3e). Based on these spatial relationships to other participants, we computed the eigenvector centrality (EC) of each participant as a measure of spatial proximity to others. Figure 3f shows the relationship between the rank EC (computed within each group) and average rewards. In random environments there was a negative trend between spatial proximity and reward (*r_τ_* = −.19, *p* = .081). In smooth environments, we find a non-linear relationship, where intermediate levels of proximity produced the highest performance. Participants with a rank EC of 2 performed better than their peers with lower EC who were the most distant from the group (*t*(20) = 2.3, *p* = .032, *d* = 1.0), but with no significant difference to peers with higher EC (*t*(31) = 1.7, *p* = . 107, *d* = 0.6), who were more central. These results suggest that an intermediate level of spatial proximity may have facilitated better performance in smooth environments, although there was no significant disadvantage to being more central. Thus, in smooth environments where social information was predictive of other rewards in the same area, it was better to be more central—despite increased competition for rewards—than to be at the outskirts, where imitation was more expensive due to increased travel costs.

#### Social Influence

In order to more directly measure social influence, we adapted methods developed to analyze the movement patterns of geo-tracked baboons in the wild (Strandburg-Peshkin, Farine, Couzin, & Crofoot, 2015). This allows us to detect discrete “pull” events over arbitrary time scales, where the movement patterns of one participant (leader) pulls in another (follower) to imitate and forage in the same vicinity (Fig. 4).

**Figure 4:**
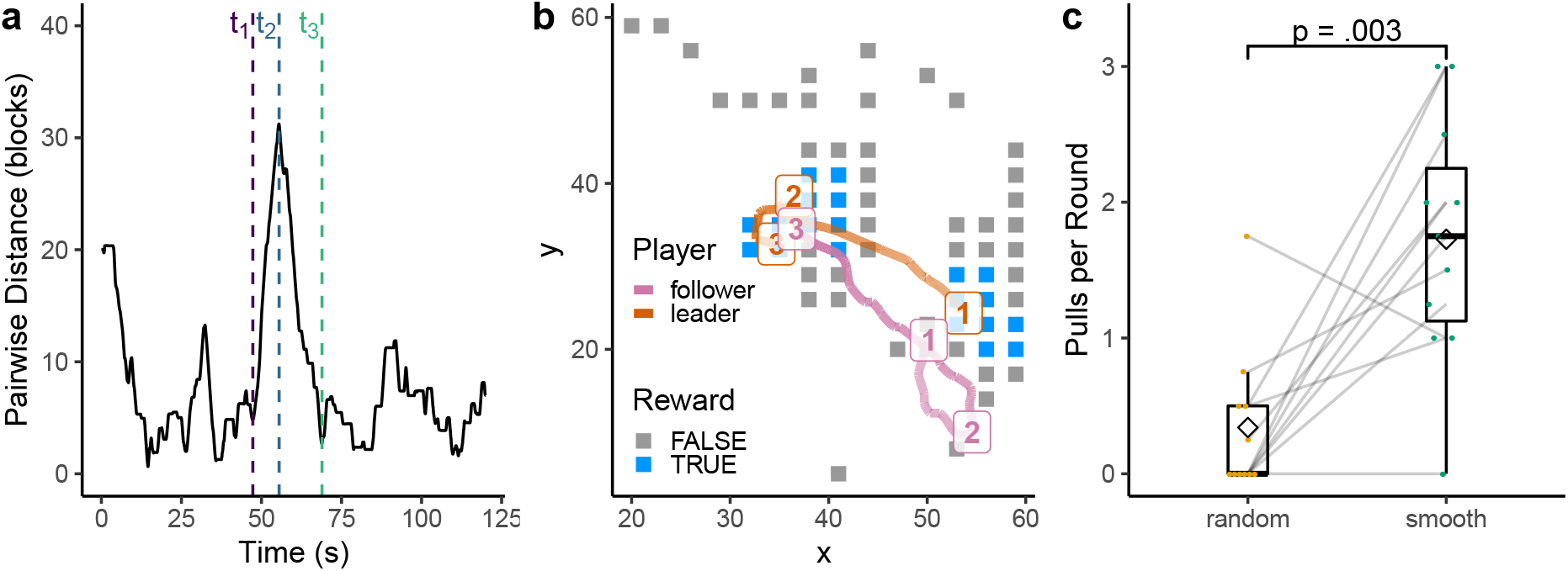
Pull events. **a**) Candidate pull events were selected from min-max-min sequences (dashed lines) of the pairwise distances between participants. These candidate sequences were then filtered by strength, disparity, leadership, and minimum duration (see text). **b**) An example of a successful pull. The orange and pink lines indicate the trajectories of the leader and follower (respectively), where the timepoints [1, 2, 3] correspond to the min-max-min sequence in panel a (dashed lines). The colored blocks illustrate the foraged blocks up until *t*_3_. **c**) The average number of successful pull events in each session (connected dots). The Tukey boxplots illustrate the aggregate statistics with diamonds showing the group, and the *p*-value is for a paired *t*-test.

We first computed the pairwise distance between all participants (Fig. 4a) and defined candidate pull events from min-max-min sequences. These candidate sequences were then filtered based on *strength, disparity, leadership*, and *duration* in order to be considered a successful pull.

**Strength** *S_i,j_* defines the absolute change in dyadic distance relative to absolute distance:

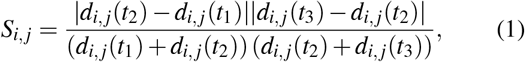

where *d_i,j_*(*t_k_*) is the dyadic distance between participants *i* and *j* at time *k* ∈ [1, 2, 3] (corresponding to the timepoints of the min-max-min sequence). We required pull events to have a minimum strength of *S_i,j_* > .1, such that they correspond to meaningful changes in spatial proximity rather than minor “jitters” at long distance.

**Disparity** δ_*i,j*_ defines the extent to which one participant moves more than the other in each segment, relative to the total distance moved by both participants:

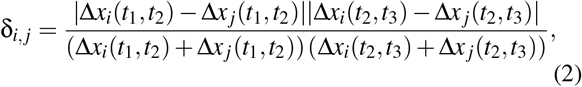

where Δ*x_i_*(*t*_1_, *t*_2_) is the displacement between *t*_1_ and *t*_2_. We filtered pull events to have a minimum disparity of δ_*i,j*_ > .1, such that changes in spatial proximity were asymmetrically driven by one of the interaction partners.

**Leadership** is a simple binary filter requiring that the participant who moved more in the first segment (*t*_1_ to *t*_2_) moved less in the second segment (*t*_2_ to *t*_3_). We refer to the participant who moved the most in the first segment max_*a*∈(*i,j*)_ Δ*x_a_*(*t*_1_, *t*_2_) as the *leader* and the participant who moved the most in the second segment max_*b*∈(*i,j*)_Δ*x_a_*(*t*_2_, *t*_3_) as the *follower*. Thus, successful pulls are defined as *a* ≠ *b*, where the leader and follower are separate participants.

**Duration** was the final filter, where we required pulls to be at least 3 seconds in duration (since it takes 2.25s to destroy a block). After all filters were applied, the average pull duration was 15s ± 0.73 (SEM).

Altogether, we detected 135 successful pull events from the group rounds in our data. Figure 4a-b shows an example of a successful pull in a smooth environment. At *t*_1_, both the leader (orange) and follower (pink) are in similar locations, but begin to move in opposite directions. At *t*_2_, the leader has found a new patch at the center of the map while the follower has been unsuccessful at the south-east corner. Between *t*_2_ and *t*_3_, the follower turns around and starts moving towards the leader and begins foraging in the same proximity.

When comparing the influence of environment on the frequency of pull events, we found a higher frequency of pulls in smooth than random environments (*t*(10) = 4.0, *p* = .003, *d* = 1.9; Fig. 4c). This effect was also amplified over successive rounds. We fit a Bayesian mixed effects Poisson regression model to predict the number of pull events in each round, using the reward environment and round number as predictors, and treating session as a random intercept. Pull events increased over rounds in smooth environments (*b* = 0.42,95% HPD: [−0.02, 0.89], *p*(*b* > 0) = .97), whereas they tended to decrease over rounds in random environments (*b* = −0.27, 95% HPD: [−0.76, 0.21], *p*(*b* < 0) = .86). This adaptation of pull frequency was consistent with the fact that social information was predictive of rewards in smooths environment, but had no value in random environments.

### Discussion

Using a collective spatial foraging experiment implemented in an immersive virtual environment, we were able to bring together an unprecedented combination of behavioral data, including spatial trajectories, visual field data, and their complex social interactions. By analyzing foraging patterns, social interactions (visual and spatial), and social influence, we were able to shine new light on how individuals in groups negotiate the balance between social and individual learning.

By manipulating the reward structure, we were able to study how individuals in groups adapt their search strategies to the value of social information. Smooth environments had predictable rewards, such that observing when another player finds a reward provided actionable information about where to search next. In contrast, random environments had no predictable pattern of rewards, and thus time spent observing other players came only at the cost of lost opportunities for individual foraging without any benefits. Accordingly, participants observed each other less in random environments, and were less susceptible to social influence, as captured by a lower frequency of pull events. In both cases, these patterns were amplified over successive rounds, suggesting a gradual rather than sudden adaptation of social learning strategy.

Furthermore, our visibility analysis indicates that groups achieved a balance between individual and social learning through specialization rather than homogeneous strategy use. In smooth environments, participants specialized as either the target of social attention or source of it. This asymmetric social attention structure may help prevent runaway information cascades. Attention selectively directed towards participants who rely more on individual learning avoids creating highly correlated social information, which is a key feature of maladaptive information cascades (Toyokawa et al., 2019; Tump et al., 2020) and also polarized echo chambers that develop through online social media networks (Baumann, Lorenz-Spreen, Sokolov, & Starnini, 2020).

The main limitation of these current results is our small sample size due to COVID-19 related restrictions on lab experiments. Even though we have incredibly rich spatial-temporal data from each participant, we have relatively weak statistical power in comparing different groups. Therefore, we have abstained from statistical analyses relating group-level characteristics (e.g., social network characteristics and pull events) to reward outcomes. For the same reason, we have not yet studied the stability of individual differences in strategy use. Recent work has found that differences in social learning strategies (imitation vs. emulation) emerge early in human ontogeny (Yu & Kushnir, 2020), which may persist as stable personality traits. Exploring the interplay between flexibility in social learning strategies, and consistent individual differences in social learning strategies is a fruitful avenue for future work.

Finally, future work using computational models and agent-based simulations will allow us to tackle a wider range of research questions and improve our understanding of the interaction dynamics that shape our social learning strategies. In sum, we have only begun to fully leverage the richness of this experimental paradigm.

## Acknowledgements

We thank Jann Wäscher and Philip Jakobs for help with running the experiments. CMW is supported by the German Federal Ministry of Education and Research (BMBF): Tübingen AI Center, FKZ: 01IS18039A and funded by the Deutsche Forschungsgemeinschaft (DFG, German Research Foundation) under Germany’s Excellence Strategy-EXC2064/1–390727645.

## Footnotes

1 The original target sample size was 128 participants, but we were forced to pause lab-based experiments since March of 2020 due to COVID-19. These current analyses thus focus on within-subject and within-session comparisons for the highest statistical power.

